# ChatGPT-Enhanced ROC Analysis (CERA): A Shiny Web Tool for Finding Optimal Cutoff in Biomarker Analysis

**DOI:** 10.1101/2023.07.13.548794

**Authors:** Melih Agraz, George Em Karniadakis

## Abstract

Diagnostic tests play a crucial role in establishing the presence of a specific disease in an individual. Receiver Operating Characteristic (ROC) curve analyses are essential tools that provide performance metrics for diagnostic tests. Accurate determination of the cutoff point in ROC curve analyses is the most critical aspect of the process. A variety of methods have been developed to find the optimal cutoffs. Although the R programming language provides a variety of package programs for conducting ROC curve analysis and determining the appropriate cutoffs, it typically needs coding skills and a substantial investment of time. Specifically, the necessity for data preprocessing and analysis can present a significant challenge, especially for individuals without coding experience. We have developed the CERA (ChatGPT-Enhanced ROC Analysis) tool, a user-friendly ROC curve analysis web tool using the shiny interface for faster and more effective analyses to solve this problem. CERA is not only user-friendly, but it also interacts with ChatGPT, which interprets the outputs. This allows for an interpreted report generated by R-Markdown to be presented to the user, enhancing the accessibility and understanding of the analysis results.

**Authors summary:** **Melih Agraz**, after graduating from Dokuz Eylül University in Izmir, Turkiye, with a major in Mathematics from the Department of Education, he pursued his Master’s degree at the same university in the field of Statistics. He furthered his education by obtaining a Ph.D. in Statistics from the Middle East Technical University, Ankara in Turkiye. As part of his academic journey, he also served as a Fulbright postdoctoral researcher at UC Berkeley in the United States. Following this, he worked as a postdoctoral researcher at Brown University and Beth Israel Hospital of Harvard Medical School. Currently, he holds the position of Assistant Professor in the Department of Statistics at Giresun University in Turkey.

**George Em Karniadakis** is Professor of Applied Mathematics and Engineering at Brown University. He is a member of the National Academy of Engineering of USA. His interests include stochastic multiscale modeling of physical and biological systems, physics-informed machine learning, and deep neural operators. He has co-authored over 500 papers and five books, and he has the highest h-index in Applied Mathematics according to Google Scholar.

## Introduction

A diagnostic test is a medical or statistical procedure that assesses an individual’s likelihood or propensity for a specific disease. In diagnostic tests with binary outcomes, performance is evaluated in terms of performance measures, sensitivity, and specificity. The Receiver Operating Characteristic (ROC) curve is a widely used visual tool in assessing the performance of diagnostic tests or classification models. It illustrates the relationship between true positive rates (TPR), also known as sensitivity, and false positive rates (FPR), which is equivalent to 1-specificity. The area under the curve (AUC) derived from the ROC curve is widely recognized as a reliable measure of accuracy. It offers meaningful interpretations regarding the performance of a diagnostic test [16]. ROC curve analysis, crucial for assessing the performance of a biomarker relative to the outcome, was first employed in the 1950s for signal detection [10]. Its inception traces back to World War II when it was used in radar technology [11]. When establishing a cutoff value, two distinct statistical approaches can be utilized: biomarker-oriented and outcome-oriented methods [18]. In diagnostic testing, a cutoff value is used to classify individuals as positive or negative based on the test results. The biomarker-oriented method determines the optimal cutoff value using specific statistics like mean, mode, or certain percentile values of the biomarkers, without considering the response variable. On the other hand, the outcome-oriented method determines the optimal cutoff value by examining the relationship between the biomarker and the response variable. This approach often yields more accurate cutoffs [20].

There are several tools available, both commercial and open-source, for calculating ROC curves and determining optimal cutoff points. Some commonly used commercial tools for this purpose include SPSS, MedCal, Stata, and Minitab. Additionally, there are powerful and freely available programming languages like R and Python that can be utilized for these tasks. R, in particular, is well-known for its extensive capabilities in statistical analyses and offers numerous packages specifically designed for calculating ROC curves and determining optimal cutoff points. These include ROCR [6], plotROC [3], pROC [13], OptimalCutpoints [7], precrec [5], and ROCit [4]. In addition, there are web-based tools powered by the shiny framework for ROC curve analysis and cutoff value calculation. For instance, easyROC is a shiny-based web tool for ROC curve analysis [8] that calculates ROC curves, cutoff points, and sample sizes. Another shiny tool, the cutoff finder [9], calculates ROC curves and cutoff values, and additionally enables users to conduct survival analysis.

Processing the data in ROC curve analysis, calculating the statistical results of the biomarker, displaying them graphically, and calculating the cutoff values by performing the ROC curve analysis is always difficult for those who are not familiar with the R programming language, especially for medical doctors. Therefore, in this study, a user interface web tool is developed for those who do not know programming languages or are looking for faster solutions in ROC curve analysis. Additionally, a key feature of this tool is its ability to automatically generate interpretations of the outputs, enhanced by ChatGPT technology. We are providing the ChatGPT-Enhanced ROC Analysis (CERA) tool which performs ROC curve analysis and optimal cutoff values with a shiny user-interface web tool.

In this paper we define the ROC curve analysis in the Method section, and we represent and discuss the output of the CERA tool in the Results section. Finally, we briefly discuss the tool and outputs in the Conclusion section.

## Materials and methods

The Receiver Operating Characteristic (ROC) curve is a graphical representation extensively utilized to illustrate the discriminatory accuracy of a diagnostic test or marker between two distinct groups. This technique has garnered significant attention owing to its effectiveness in showcasing the marker’s discriminatory accuracy. Essentially, the ROC curve offers a visual framework for comprehending the proficiency of a specific marker in distinguishing between two disparate populations.

Diagnostic tests categorize individuals into two groups, such as positive/negative, and there is always a potential for error, given that these two groups may overlap. Let us assume an example that two groups, positive and negative, overlap as depicted in Figure 1b, and these groups are distinguished by the threshold *c* as shown in Figure 1a; we need to elucidate the types of errors that might occur. As can be observed in Figure 1b, values greater than the *c* cutoff are predicted to be positive (*T* +), while smaller values are predicted to be negative (*T−*). However, the region we denote as FP (Type I error) is actually negative, but it is predicted as positive. The remaining predictions, which are correctly identified as positive, are denoted as TP. Similarly, values smaller than *c* are correctly predicted as negative (TN), while incorrect predictions are labeled as FN (Type II error). If we can generate tables (as shown in Figure 1a) for each candidate threshold, and calculate the sensitivity versus (1-specificity) values, we can create the figure shown in 1c. This figure displays the ROC curve for two overlapping groups. As indicated in Figure 1c, the diagnostic test appears to be a more effective discriminator than a random guess.

**Fig 1.**
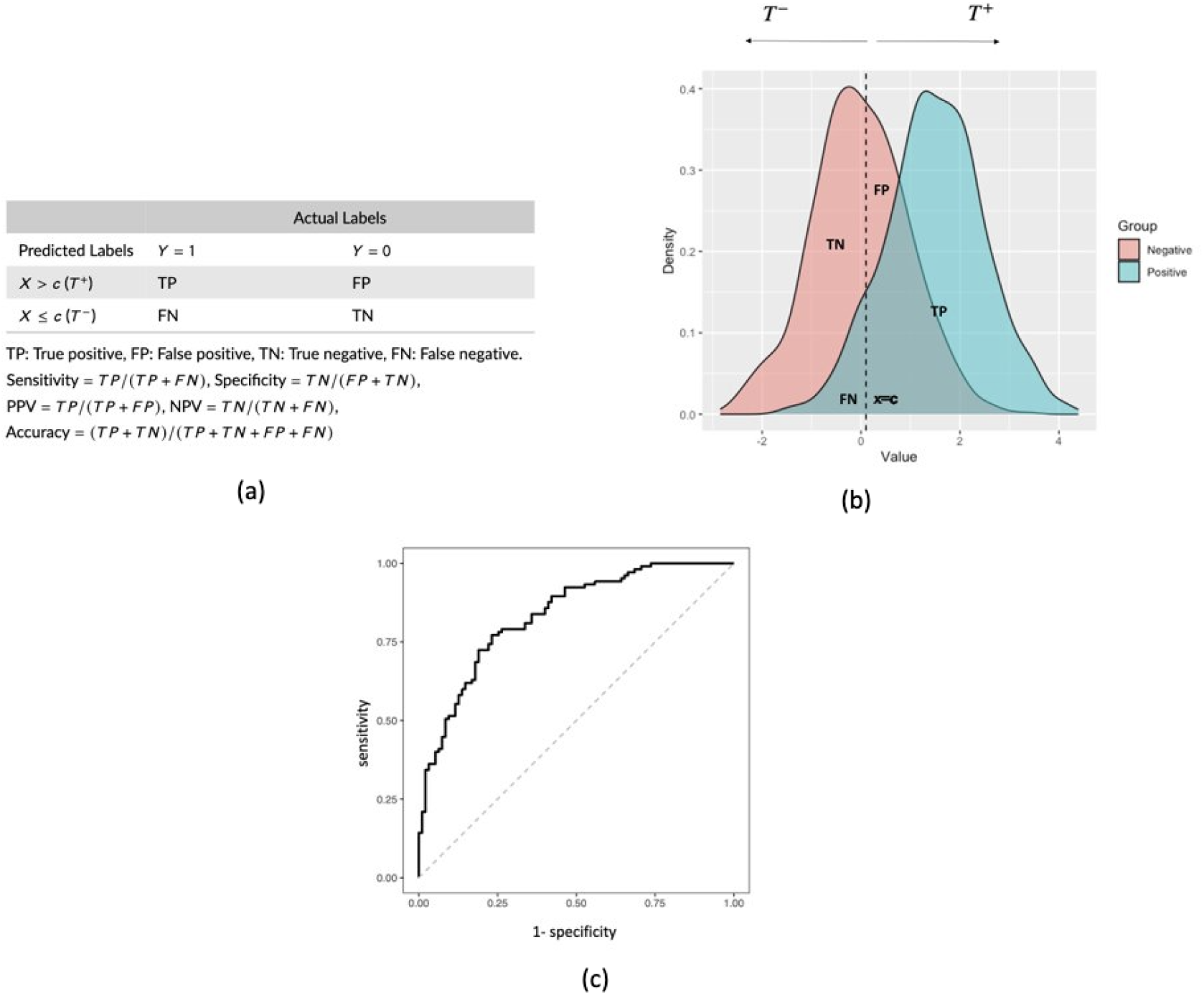
Overview of the ROC analysis process and performance metrics. (a) 2×2 classification table illustrating the performance of the classifier. The table demonstrates the different types of predictions made by the classifier based on the chosen cutoff value, including true positives (TP), false positives (FP), true negatives (TN), and false negatives (FN). (b) Hypothetical distributions of decision-making, demonstrating overlap between the positive and negative labels. The overlapping regions represent the potential for misclassification by the classifier. (c) ROC curve created from two overlapping distributions as in Figure (b), highlighting the performance of the classifier at various cutoff points. The ROC curve gives a visual depiction of the classifier’s discriminating power by demonstrating the trade-off between sensitivity and specificity for various threshold values.

We are using the shiny tool to convert the R-generated ROC curve analysis to a user interface web tool. A typical shiny application is composed of two main elements represented in Figure 2: (i) **user interface (UI)** that constructs the visual appearance, and (ii) **server** that contains instructions for executing and updating the objects presented in the UI. In addition to that, the proposed CERA tool has a ChatGPT API connection as represented in Figure 2.

**Fig 2.**
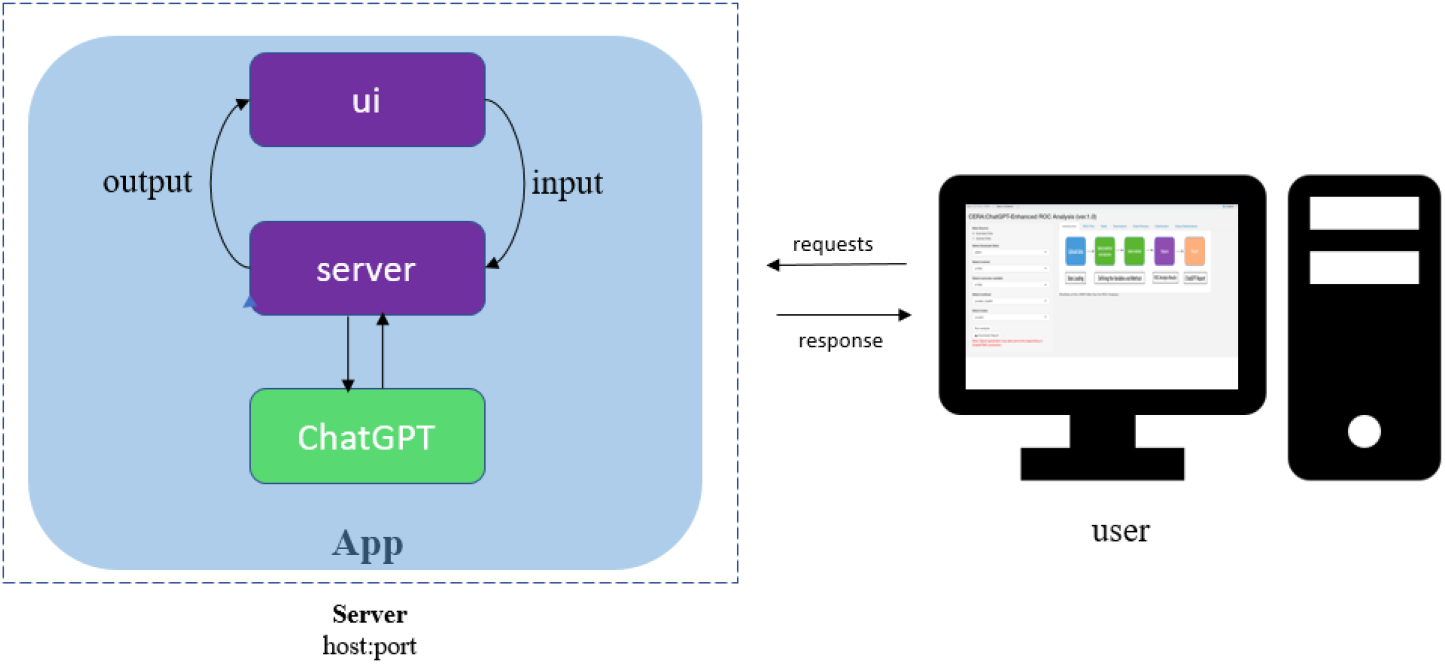
A Shiny Workflow Diagram Illustrating the UI, Server, and ChatGPT API Connection. The diagram visually represents a dynamic workflow demonstrating the interactions between the user interface (UI), the server, and the ChatGPT API within the context of data analysis and reporting. The CERA tool enables users to upload their data, select markers, and choose appropriate methods for analysis. This step signifies the user’s input and customization of the analysis process. Once the necessary inputs are provided, the user can proceed by clicking on the ”Run analysis” button. This action triggers the tool to perform the ROC analysis. Upon completion, the user can proceed by clicking on ”Download Report” to establish a connection with the ChatGPT API. This step allows the user to receive a report generated by ChatGPT, which includes interpretations of plots and tables.

In the shinny app, the UI determines the visual appearance and layout of the web application. The UI code defines the organization of the application, including the various input elements (e.g., sliders, text boxes) and output components (e.g., charts, tables) that users interact with. On the other hand, the server is responsible for generating dynamic outputs based on user inputs and updating them in real-time. It handles the computational and data processing tasks of the application. When the user interacts with the UI, the server processes the input data, performs the necessary calculations or manipulations, and sends the computed results back to the UI for display. In this study, we have incorporated the ChatGPT API connection using the gptchatter [25] library to provide interpretations for the ROC curve analysis results. The default GPT-3.5 version is utilized by this library. The connection to the API is established when the user clicks on the ”Download Report” button, and the report is generated based on the results of the ROC curve analysis. The report consists of three sections: Introduction, Results (with subsections for data quality checking, ROC plot, and performance measures), and Conclusion. The API connection prompts are fixed and not user-modifiable. The CERA tool establishes an API connection for each section and subsection of the report, resulting in a total of five API connections for each report preparation.

### ChatGPT-Enhanced ROC Analysis (CERA) web tool

The proposed CERA web tool, freely available at https://datascicence.shinyapps.io/ROCGPT/, is developed using the shiny app [15] and pROC [13] R package. It calculates optimized cutoff values for the *X* continuous variable (biomarker) and the *Y* binary outcome variable. In addition to providing the ROC curve plot and the distribution of the biomarker, it also shows statistical results such as mean, mode, median, and standard deviation of the biomarker. It also visualizes the relationship between the biomarker and the binary response variable with a boxplot class distribution graph. Finally, CERA can report this information in three parts with ChatGPT-supported interpretations: Introduction, Results and Conclusion sections with Data Quality Checking, ROC Plot and Perfromance Measures subsections in Results section. The reporting is connected and data is retrieved via the ChatGPT, developed by OpenAI [21]. This data is converted into a Rmarkdown HTML format that can be downloaded by the user.

### Workflow steps

The workflow involved in data processing, as illustrated in Figure 3, is listed in the subsequent items:

**Fig 3.**
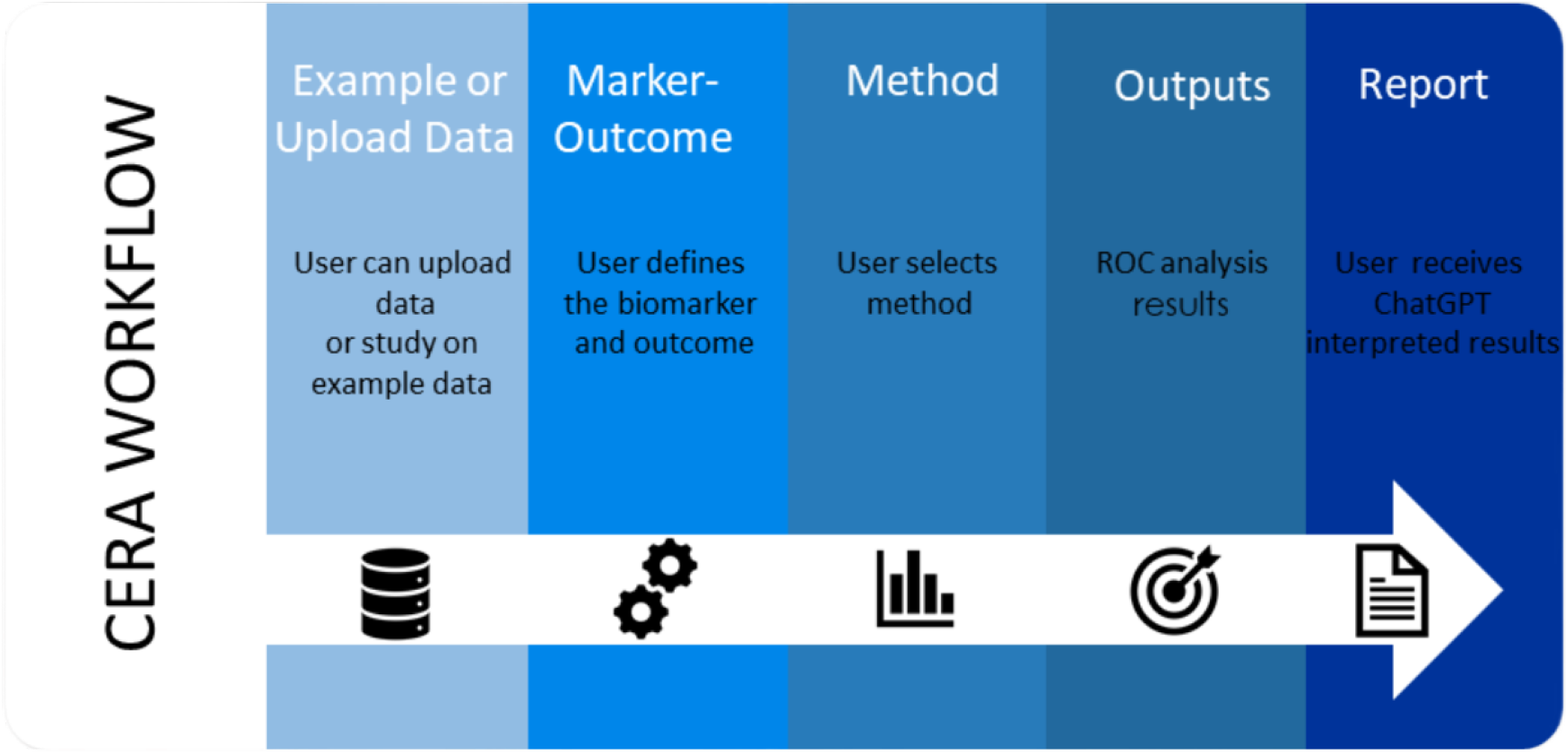
Workflow of the CERA Web Tool for ROC Analysis. The CERA workflow consists of the following steps: Example or Upload Data: In this step, the user can upload their own data or example datasets provided by CERA tool to the system. Marker-Outcome: In this step the should select the biomarker and the outcome variable. Method: The user selects the method they want to apply in the study. Outputs: This is the step where ROC graph and performance characteristics are taken. Report: By clicking ”Download Reprt”, the user can download the report prepared with ChatGPT under the sections of Intorduction, Results and Conclusion of the CERA outputs.

1. The user can upload data in a comma-separated format, with the header first row. The user also has the option to work with example datasets provided in the tool, such as the aneurysmal subarachnoid hemorrhage (aSAH) dataset [14], the PIMA Indian dataset [24], the Wisconsin breast cancer dataset [23], or simulated non-alcoholic fatty liver disease (NAFLD) data.
2. The user defines the biomarker and binary response from the uploaded data.
3. The user selects the cutoff method from the Select Method part. The Select Method part has youden_topleft, cutoff, maximized and min_pvalue_approach.
  - youden_topleft: Users can select either the Youden Index (youden) method or the Closest Top-Left (closest.topleft) method under the ”Select Index” conditional panel.
  - cutoff: Users can specify a specific cutoff value.
  - maximized: Users can select the minimum and maximum values for specificity and sensitivity under the ”Select Constrain Metric” conditional panel.
  - min_pvalue_approach: Users can select the minimum p-value approach.
4. The CERA tool analyses the data, generates ROC curve plots and related results.
5. There is a ”Download Report” button to generate a ChatGPT generated report and receive the output in an HTML file.

The cutoff methods used in the tool can be determined as follows.

- Youden’s Index: Youden’s Index was first introduced by William John Youden [1] to optimize the cutoff value of the ROC curve. This optimization is done by maximizing the difference between sensitivity and (1-specificity), as represented by Youden’s statistics in the Equation 1.

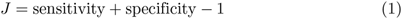

The range of Youden’s index is from -1 to 1. A value of 0 suggests that the performance of the classifier is equivalent to a random selection, whereas a value of 1 means perfect classification performance. As the value of J increases, the performance is deemed better, given that it implies a greater disparity between the true positive rate and the false positive rate. Youden’s index has a simple and intuitive interpretation: it corresponds to the point on the curve that is farthest from the ”chance” line [17].
- Closest top-left: This method chooses the best cutoff threshold by identifying the value closest to the upper-left corner of the ROC curve as shown in Equation 2, to find the optimum cutoff in ROC curve analysis. This method calculates the optimal cutoff by the min value of the square distance between (0,1) and the value on the ROC curve, i.e.

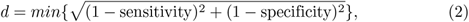

where *d* is calculated for each cutoff, and the cutoff that gives the minimum distances is selected as the optimal cutoff value. It is important to note that the top-left method, being a simple heuristic, may not consistently provide the optimal threshold for a given problem.
- Minimum p-value approach: Cutoff values are calculated by minimum p-value approach by creating a 2 × 2 table for each potential cutoff value and selecting the optimal cutoff, *c*, that maximizes the chi-square statistics or minimizes the *p* value. A disadvantage of the min-*p*-value approach is that it can result in a high number of Type I errors.
- Minimum sensitivity/specificity: The user can define minimum or maximum specificity or sensitivity values, and thus the defined value takes the minimum of that value. For example, if the user defines a value of 0.7 for specificity, the performance measure results show the value with a minimum specificity of 0.7. This method works on *cutpointr* function [19].
- Biomarker-oriented approaches: The tool automatically calculates the mean, median, and mode. These results can also be used as cutoff points in biomarker-oriented approaches.

In addition, the user can manually define the cutoff value and see the performance measures of it.

## Results

In order to demonstrate the effectiveness of our CERA tool, we use four example distinct datasets: i) the aneurysmal subarachnoid hemorrhage (aSAH) dataset [14], which is available in pROC package [13], ii) simulated NAFLD dataset, iii) PIMA Indian dataset [24], and iv) Wisconsin breast cancer dataset [23]. However, for the purpose of demonstrating the effectiveness of the CERA tool, our focus will be on presenting and discussing the results obtained from the aneurysmal subarachnoid hemorrhage (aSAH) dataset exclusively. This dataset will serve as a representative example to showcase the capabilities and performance of the CERA tool in analyzing and interpreting the ROC curve analysis results.

### Aneurysmal subarachnoid hemorrhage (aSAH) data

This data [14] comprises 113 individuals with aneurysmal subarachnoid hemorrhage including 6 explanatory variables and 1 binary outcome variable (Good/Bad). The explanatory variables are Glasgow outcome score at 6 months (GOS6), gender, age, world federation of neurological surgeons (WFNS), S100*β*, and NDKA. However, we retain only the S100*β* variable, considering it as a biomarker for the purpose of ROC curve analysis.

### Non-alcoholic fatty liver disease (NAFLD) simulated data

Nonalcoholic fatty liver disease (NAFLD), recognized as the most common cause of liver disease [12], is frequently assessed using the hepatic steatosis index (HSI) as a screening tool [2]. Consequently, we simulate a dataset for NAFLD, consisting of 300 patients, with a binary response variable (NAFLD/non-NAFLD, labeled as 1/0). In this simulated dataset, data labeled ’1’ (indicating NAFLD) has a mean HSI of 55 with a standard deviation of 10. Conversely, data labeled ’0’ (indicating non-NAFLD) has a mean HSI of 40, also with a standard deviation of 10.

### PIMA Indian dataset

Another example dataset available in the CERA tool is the PIMA Indian dataset sourced from Kaggle [22]. This dataset is particularly relevant as it shows the high prevalence of type 2 diabetes among women of PIMA Indian heritage who are at least 21 years old. It consists of eight variables and a target variable, with a total of 768 observations. The features included in the dataset are as follows: pregnancies (number of times pregnant), glucose (plasma glucose concentration), blood pressure (diastolic blood pressure, mm Hg), sin thickness (triceps skin fold thickness in mm), insulin (mu U/ml), BMI, diabetes pedigree function, and Age (years). The target variable, namely Diabetes, for binary classification takes the values of 0 or 1, where 0 indicates a negative test result for diabetes, and 1 indicates a positive test result.

### Wisconsin breast cancer dataset

The Wisconsin breast cancer dataset [23] consists of 30 features along with the target variable ”Breast Cancer”. This dataset contains a total of 569 observations. The target variable represents the presence of breast cancer and has two classes: malignant (M) and benign (B). Among the observations, there are 212 cases classified as malignant and 357 cases classified as benign. The dataset provides valuable information for conducting analysis and exploring the relationship between the features and the presence of breast cancer.

The user can use the provided example datasets or have the option to upload their own data separated by commas. It is important to note that user-uploaded data must contain a header row with variable names. This data can be uploaded by selecting the Upload Data radio button as shown in Figure 4.

**Fig 4.**
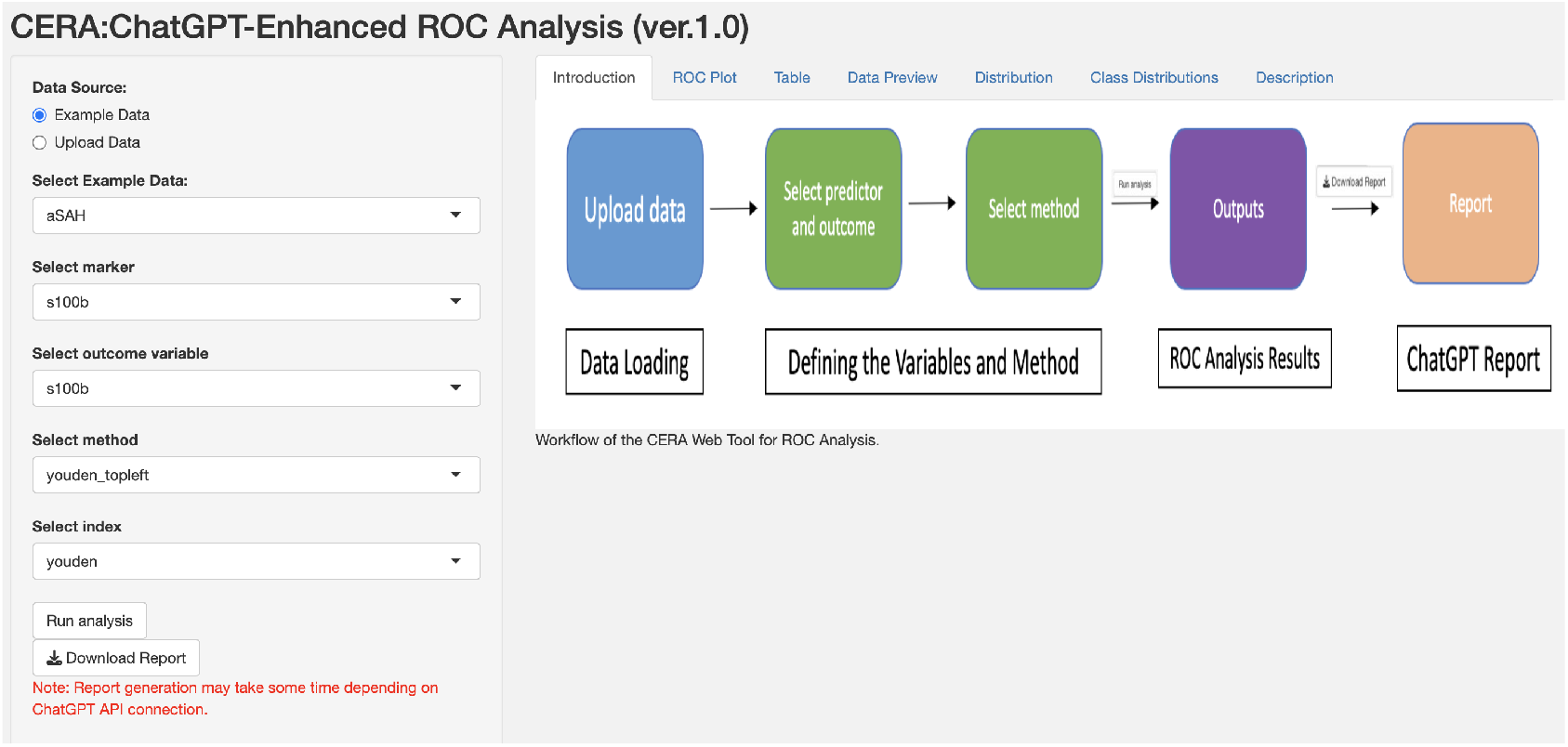
Uploading Data and Using Example Datasets in the CERA Tool. The figure shows the process of uploading data and using example datasets within the CERA tool. The tool has four example datasets: three publicly available (PIMA, aSAH, and Wisconsin breast cancer data) and one simulated dataset (NAFLD) derived from real data. Users have the option to select one of these example datasets or upload their own data. For uploaded data, it should be in comma-separated format with the first row serving as the header containing variable names.

**Fig 5.**
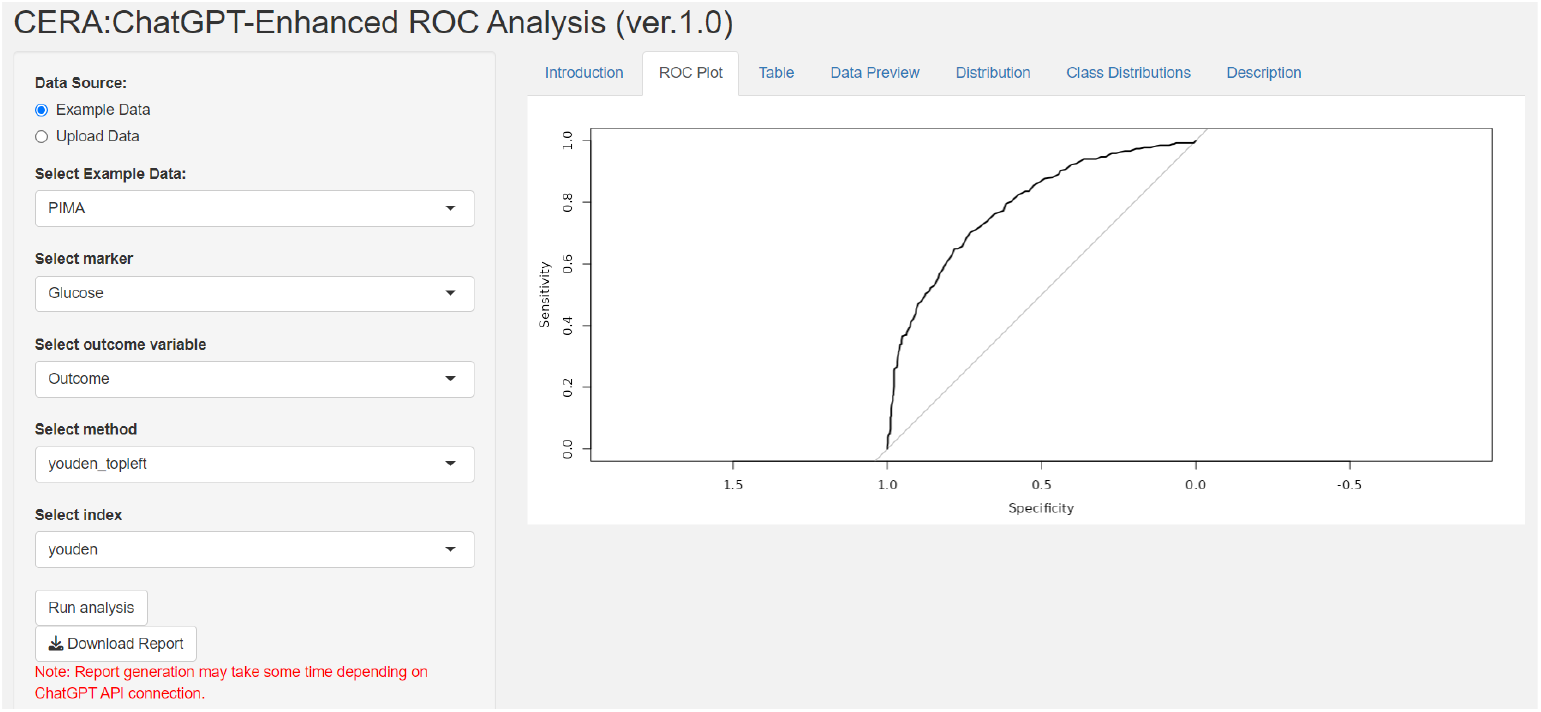
ROC plot output of the CERA tool. The figure shows a ROC plot, illustrating the relationship between specificity and sensitivity. A classification model’s or classifier’s effectiveness is graphically depicted by the ROC curve. In this example, the ROC curve demonstrates the Area Under the Curve (AUC) value will be higher than 0.5. While the figure provides a visual representation of the ROC curve, users can refer to the Tables panel within the CERA tool to access specific AUC values.

After the data is uploaded, the user can start the analysis by clicking on ”Run analysis” button. The CERA tool then automatically analyses the data, presenting the results in six different Tab panels. These panels are:

- ROC Plot: This panel brings the ROC curve of the analysis.
- Table: This panel provides performance measures of the output such as cutoff value, sensitivity, specificity, positive predictive value (PPV), negative predictive value (NPV), accuracy, AUC, lower bound of the AUC confidence interval, and upper bound of the AUC confidence interval.
- Description: This panel offers brief information about the method used.
- Distribution: This panel shows the distribution of the biomarker, along with statistical analysis of the biomarker such as mean, median, and standard deviation.
- Class Distribution: This panel provides a boxplot of the labels of biomarkers.

We illustrate the capabilities of the CERA tool by analyzing the aSAH data [14]. Users can first choose the optimal cutoff method from the options provided under ”Select method”. As detailed in the Method section, this feature offers four different cutoff methods and an option to manually select a cutoff value. If the user wishes to focus on a biomarker-oriented approach, the CERA tool also provides biomarker statistics under the ”Distribution” section. Once the user defines the biomarker and outcome variables and selects a cutoff method, they can initiate the analysis. The tool generates the results after the user initiates the ”Run analysis” function.

Figure 6 presents the distribution classes of the biomarker, while Table 1 provides the performance measures as depicted in the ‘Table’ panel of the CERA tool. The tool calculates a threshold of 0.20 and an AUC of 0.73 (with confidence intervals ranging from confidence interval lower boundary (CI:Lower B.) 0.63 to confidence interval upper boundary (CI:Upper B.) 0.83). Additionally, it computes sensitivity, specificity, positive predictive value (PPV), negative predictive value (NPV), and accuracy. The ‘Description’ panel offers explanations for the selected methods. These definitions are fixed and are not generated by ChatGPT.

**Table 1.** Performance Measures of ROC Curve Analysis. The table represents the performance measures of ROC curve analysis, as shown in the Table Tab panel of the CERA tool. The results indicate the following metrics: threshold: 0.20, Sensitivity: 0.63, Specificity: 0.81, Positive Predictive Value (PPV): 0.65, Negative Predictive Value (NPV): 0.79, Accuracy: 0.74, Area Under the Curve (AUC): 0.73, Lower Boundary (Lower B.) of AUC: 0.63, and Upper Boundary (Upper B.) of AUC: 0.83.These performance measures provide valuable insights into the accuracy and effectiveness of the ROC curve analysis performed by the CERA tool.

| threshold | Sensitivity | Specificity | PPV | NPV | Accuracy | AUC | CI: Lower B. | CI: Upper B. |
| --- | --- | --- | --- | --- | --- | --- | --- | --- |
| 0.20 | 0.63 | 0.81 | 0.65 | 0.79 | 0.74 | 0.73 | 0.63 | 0.83 |

**Fig 6.**
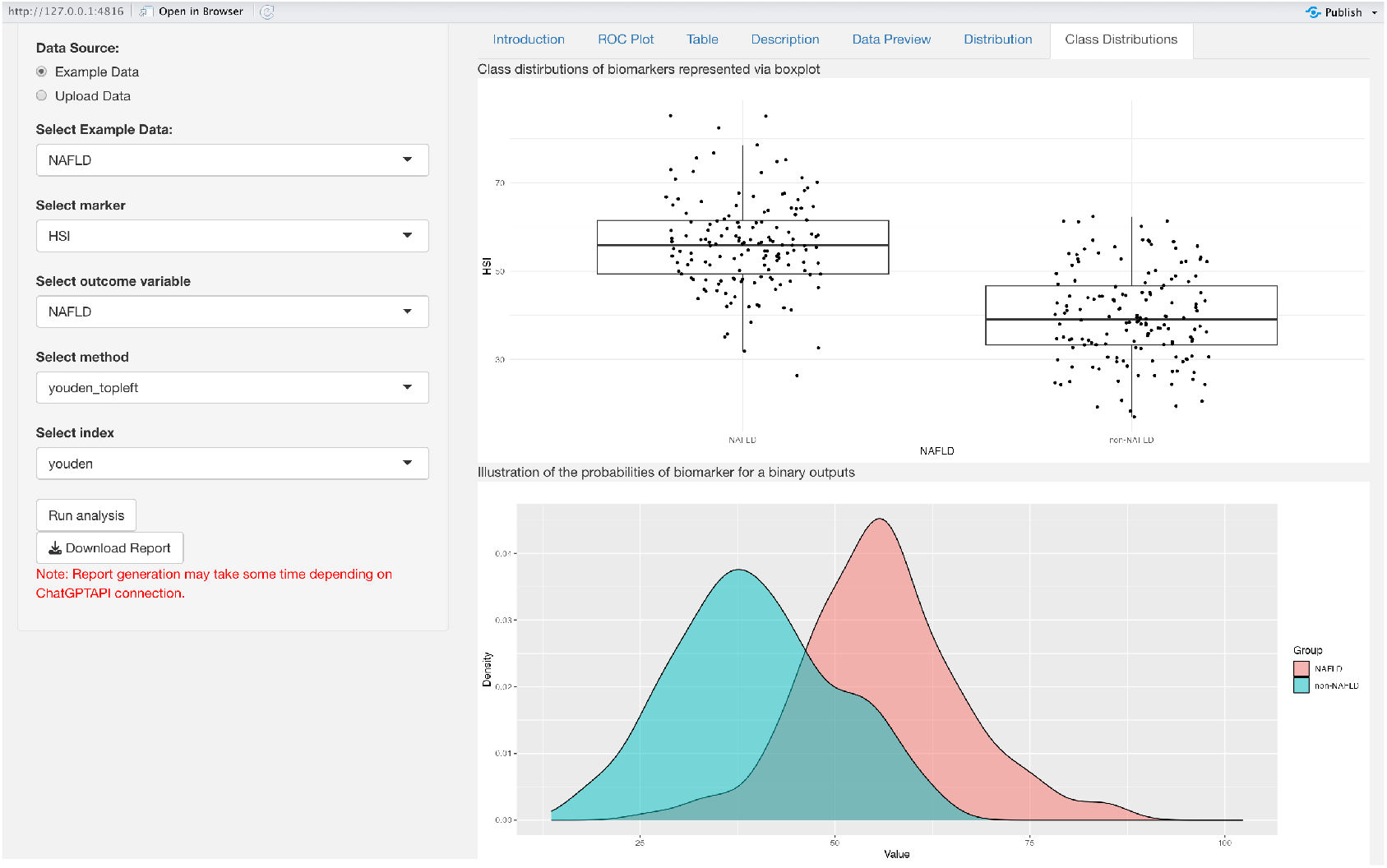
Class Distributions of Biomarker. The figure illustrates the class distributions of biomarkers using boxplots. In the top figure, the boxplot displays the class distributions of biomarkers. In the example of NAFLD, the mean value is around 50, while the non-NAFLD value is around 40. This indicates a noticeable difference between the NAFLD and non-NAFLD classes. In the bottom figure, users can obtain probability distributions of the biomarker for a binary outcome. In the example, it is observed that the NAFLD and non-NAFLD classes exhibit some degree of overlap.

The ‘Data Preview’ tab panel lists the first and last four rows of the dataset. In addition, users can access the number of observations, number of labels in the output, and missing data percentages under the following titles ”Number of Observations”, ”Number of Labels”, ”Percentage of Missing Data (%)”, respectively. With the ”Distribution” panel, statistics such as mean, standard deviation (SD) and median and biomarker distribution can be found. Finally, the ‘Class Distribution’ panel provides a boxplot generating the biomarker distributions for each label.

After generating results using the CERA tool, users may require a comprehensive interpretation and report. To facilitate this, we have included a ‘Download Report’ tab that produces a report enriched by ChatGPT, as shown in Figure 7. The generation of the report could take up to 2 minutes, depending on the ChatGPT API connection; when it is ready, the user can find it in their Download folder. We are working on optimizing this process to make it more efficient and faster. This report is divided into three sections: Introduction, Results, and Conclusion, each of which is prepared via the ChatGPT API connection [21], with the exception of the first paragraph in the Results section. This paragraph mirrors the content found in the ‘Description’ tab panel.

**Fig 7.**
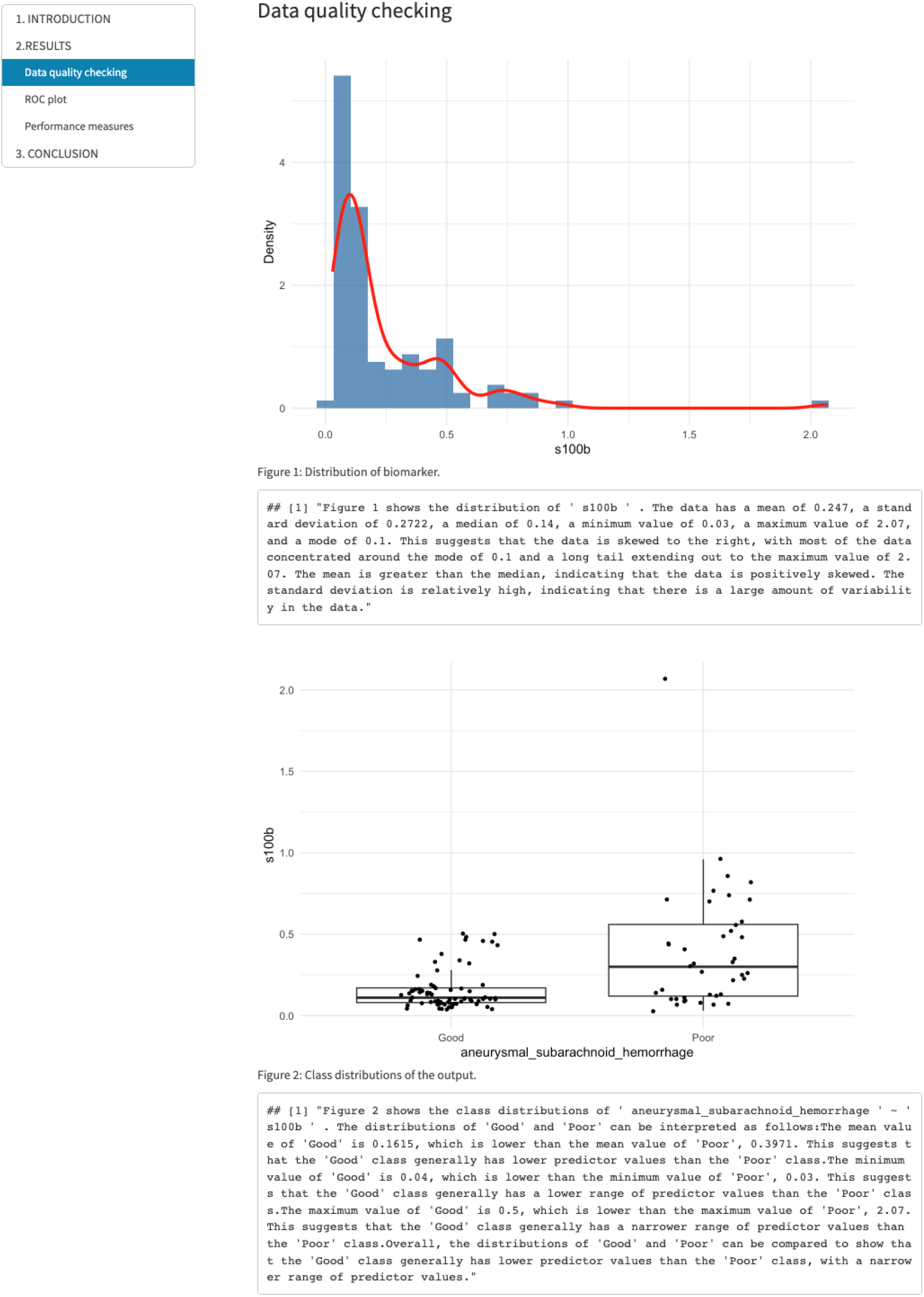
Report Generated by CERA Tool via ChatGPT -Data Quality Checking Subsection of RESULTS Section. The figure displays a subsection titled ”Data Quality Checking” within the RESULTS section of the report generated by the CERA tool using ChatGPT. This subsection includes figures from the Class Distribution panel, accompanied by interpreted results from ChatGPT. In the top figure, the distribution of the biomarker is illustrated, with a red distribution curve overlaid to provide additional information. This visualization aims to understand the distribution pattern of biomarkers. The bottom figure presents the class distributions of the output variable in the form of a boxplot that updates as ChatGPT interprets the results. According to this figure, aneurysmal subarachnoid hemorrhage data with binary class (Poor/Good) is visualized. The average value of s100b in the Poor-grade aneurysmal subarachnoid hemorrhage class is higher than the mean value of s100b in the Good-graded aneurysmal subarachnoid hemorrhage class.

- In the Introduction section, it is strongly advised that users utilize appropriate and descriptive column names for the biomarker and outcome variables instead of abbreviations. This is crucial because the column names are used as input by ChatGPT to generate interpretations. By using clear and informative column names, it enhances the accuracy and understanding of the generated interpretations.
- The Results section contains three subsections: Data Quality Checking, ROC Plot, and Performance Measures. The Data Quality Checking subsection includes the distribution plot generated in the ‘Distribution’ tab panel, which is interpreted based on the statistics of the biomarker. Additionally, the ‘Class Distribution’ tab panel plot and its interpretation are included. In the ROC Plot subsection, users can find the ROC curve generated in the ’ROC Plot’ tab panel along with its interpretation. The Performance Measures subsection features the results from the ’Table’ tab panel and their interpretations.
- Finally, the Conclusion section concisely explains the outputs and summarizes the results.

## Conclusion

ROC curve analysis is an important tool in determining the performance of biomarkers in diagnostic tests. In this study, we introduce a web-based shiny tool called CERA (ChatGPT-Enhanced ROC Analysis), designed to facilitate ROC curve analysis available at https://datascicence.shinyapps.io/ROCGPT/. This tool allows users to easily perform ROC analysis, as well as interprets and reports findings to the user via ChatGPT in the Introduction, Results and Conclusion sections. In the CERA tool, users have the option to upload their own data or use one of the provided datasets. After the data is uploaded, users define the biomarker and can choose from various optimization methods. The tool generates ROC curve graph and optimized cutoff value, as well as additional performance metrics. It also provides various statistical results regarding the biomarker. In the final step, users can generate an interpreted report with ChatGPT.

ROC curve analysis can be performed using various tools such as SPSS, Minitab or Stata or programming languages such as R and Python. However, in this work, we introduce an open source, freely accessible web tool developed with shiny. This tool has a user-friendly interface and provides many statistics necessary for data analysis, including ROC analysis. The feature that distinguishes this tool from others is that it can interpret outputs via ChatGPT and can generate detailed reports that users can use in their work.

It is important to note that the CERA tool does not store user data in any database and we cannot guarantee data security. The tool only works through the hosting platform powered by shiny.io. In future work, we plan to expand the capabilities of CERA to support multiple biomarkers. We also aim to include additional data preprocessing steps such as missing data imputation or outlier detection. We believe these improvements will enable this tool to serve as a more effective platform for data science purposes. We intend to use our own hosting platform under a different website name.

